# Recasting the cancer stem cell hypothesis: unification using a continuum model of microenvironmental forces

**DOI:** 10.1101/169615

**Authors:** Jacob G. Scott, Andrew Dhawan, Anita Hjelmeland, Justin Lathia, Anastasia Chumakova, Masahiro Hitomi, Alexander G. Fletcher, Philip K. Maini, Alexander R. A. Anderson

**Affiliations:** Departments of Translational Hematology and Oncology Research and Radiation Oncology, Cleveland Clinic, Cleveland, OH, USA; Wolfson Centre for Mathematical Biology, Mathematical Institute, University of Oxford, Oxford, UK; Department of Oncology, University of Oxford, Oxford, UK; Department of Cell, Developmental and Integrative Biology, University of Alabama, Birmingham, AL, US; Department of Cellular and Molecular Medicine, Lerner Research Institute, Cleveland Clinic, Cleveland, OH; School of Mathematics and Statistics, University of Sheffield, Sheffield, UK; Bateson Centre, University of Sheffield, Sheffield, UK; Department of Integrated Mathematical Oncology, Moffitt Cancer Center and Research Institute, Tampa, FL, USA

## Abstract

Since the first evidence for cancer stem cells in leukemia, experimentalists have sought to identify tumorigenic subpopulations in solid tumors. In parallel, scientists have argued over the implications of the existence of this subpopulation. On one side, the cancer stem cell hypothesis posits that a small subset of cells within a tumor are responsible for tumorigenesis and are capable of recapitulating the entire tumor on their own. Under this hypothesis, a tumor may be conceptualized as a series of coupled compartments, representing populations of progressively differentiated cell types, starting from stem cells. The allure of this model is that it elegantly explains our therapeutic failures: we have been targeting the wrong cells. Alternatively, the stochastic model states that all cells in a tumor can have stem-like properties, and have an equally small capability of forming a tumor. As tumors are, by nature, heterogeneous, there is ample evidence to support both hypotheses. We propose a mechanistic mathematical description that integrates these two theories, settling the dissonance between the schools of thought and providing a road map for integrating disparate experimental results into a single theoretical framework. We present experimental results from clonogenic assays that demonstrate the importance of defining this novel formulation, and the clarity that is provided when interpreting these results through the lens of this formulation.

## Introduction

Although posited to exist over forty years ago^1^, cancer stem cells (CSCs) were first identified in 1997 by Bonnet and Dick in leukemia^2^. Since this discovery, CSCs have been shown to exist in many solid tumor types, including colon^3^, brain^4^, breast^5^ and melanoma^6^. The cancer stem cell hypothesis (CSCH) states that each tumor is composed of a cellular hierarchy, at the top of which is a population of ‘stem cells’ able to self-renew and give rise to the entire diversity of cells within the tumor. The alternate, proliferative hypothesis suggests instead that each cell in the tumor has some low level of clonogenic potential, and proliferation is driven by stochastic genetic alterations. These two models are schematised in Figure 1. The CSCH provides a framework by which to understand many different aspects of cancer progression that the proliferative hypothesis cannot explain, including: functional heterogeneity despite identical genetic states^7, 8^; resistance to chemotherapy^9–11^ and radiotherapy^12–14^; recurrence^15^; and metastasis^16^. However, the CSCH has been the subject of continual debate and modification in an attempt to maintain compatibility with experimental observations. Most importantly, there is still no consensus as to how to identify a CSC^17^. The most accepted paradigm is the use of specific cell surface markers for enrichment with propagation in specific growth conditions and functional characterization with the clonogenic or sphere-forming assay and tumorigenic potential^18^. Adding to the challenge, the number of ‘stem cell markers’ is legion, and the meaning^19^, not to mention permanence^20^, of each is itself a source of ongoing debate.

**Figure 1.**
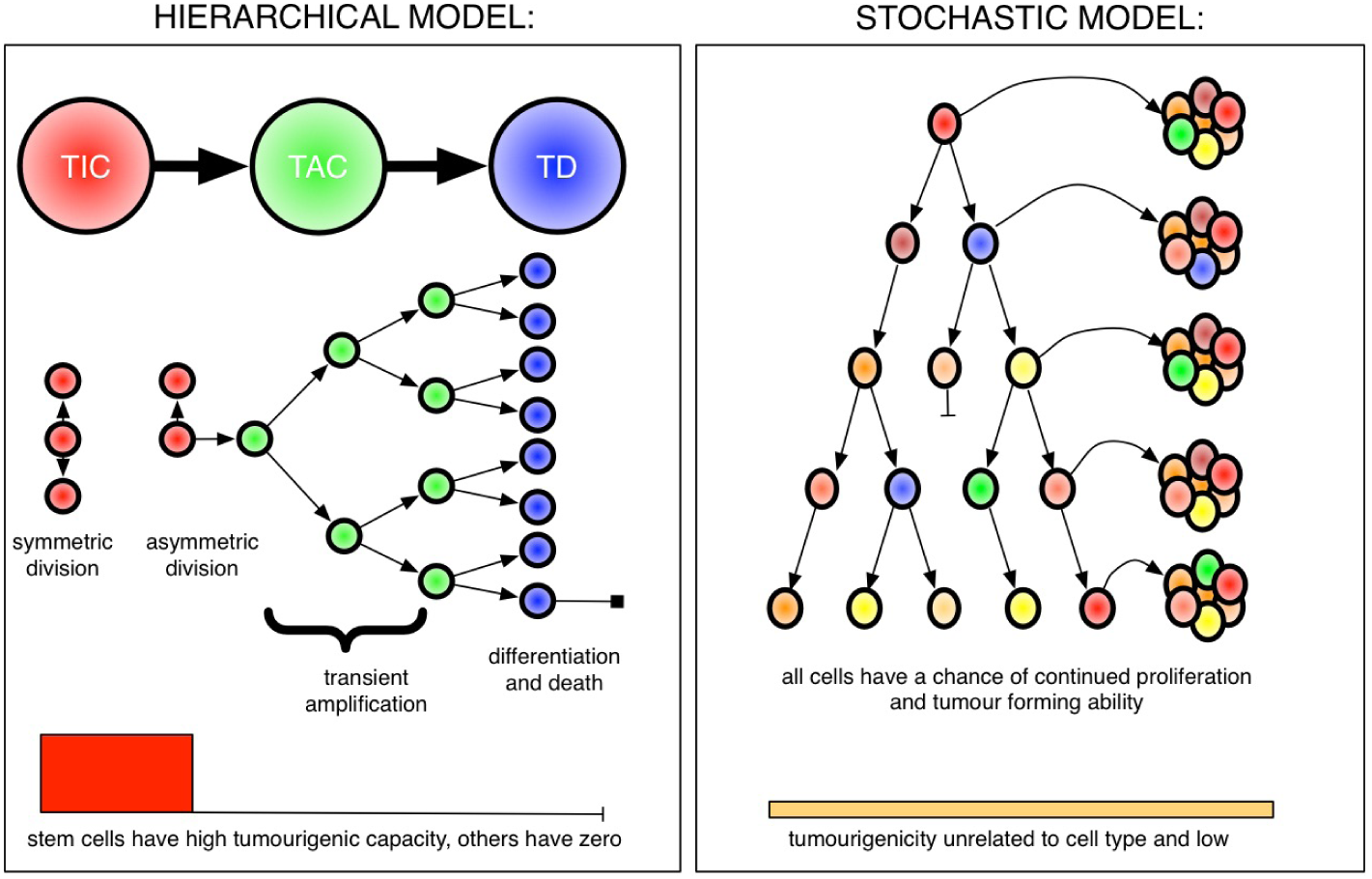
The two competing models of tumor growth regarding replicative and tumor forming potential. Left: the hierarchical, or cancer stem cell, model in which only a subset of cells, the putative cancer stem cells, have the ability to proliferate indefinitely and can recapitulate the entire tumor, while all others are doomed to differentiation and eventual death; and, Right: The standard proliferative model, in which each cell has the same ability, with low probability, to form tumors. We represent clonogenicity on the bottom of each panel where, on the left the red box means that the TICs have high probability and all others have none, and on the right that all cells have equal, low clonogenic potential.

While the CSCH has been able to explain many important aspects of cancer that the standard proliferative hypothesis has not, the arguments about its specific form continue to plague the cancer research community. In each rigorous formulation of this model, experimental data differs, and does not fit the model with the expected precision^21^. In each subsequent iteration, small adjustments to the standard hierarchical stem cell model are made. Recently, plasticity has been added to the model^22, 23^, by allowing transient amplifying cells (TACs) to dedifferentiate into stem cells to account for the experimental observations of such transformations appearing to occur randomly^24^, or in response to radiation treatment^25^ and hypoxia^26, 27^. While these modifications represent steps forward in our quest to rigorously describe, and thereby understand, cancer, they are also modifications of a model that may not be able to wholly capture the dynamics of this enigmatic disease (for an excellent review of mathematical models of the CSCH, see Michor^28^). This sort of growing dissonance is not new in science; indeed, it seems to be a conserved motif. Many examples exist where a model, like the celestial spheres in astronomy, is incrementally modified to encompass data that were not available or considered when the model was conceived. The model can become increasingly unwieldy until a new, simpler model can be postulated - as Newton’s laws explained Kepler’s observed patterns - whereupon, the cycle begins anew.

We submit that in cancer research we find ourselves in a similar situation concerning the CSCH, and that to make further progress, we must tear down the standard hierarchical architecture of the CSCH and recast it entirely. Therefore, we present a novel model of cancer cell differentiation that does not take the standard compartmental form. We instead posit a continuum model of differentiation and clonogenic state, mathematically similar to those of Hoffman et al.^29^ and Doumic et al.^30^, but composed of cells whose behaviour can change in response to environmental factors, which we model as ‘forces’. This model allows for integration of the proliferative model and the CSCH and has the potential to reconcile previous issues over surface markers, which themselves have continuous expression values, as the distinction between ‘stem’ or ‘non-stem’ cells is no longer requisite. The model also provides a new way to define a tumor’s cellular composition and progression as a dynamic distribution, and is supported by a number of recent biological observations^10, 25, 26, 31–37^ into a single modeling framework.

### Existing modifications to the canonical model cannot capture dynamic heterogeneity

The CSCH has been typically represented by a compartmental model of differentiation in which a stem cell, upon cell division, becomes a TAC, which may divide a fixed number of times creating exact copies of itself, as a ‘progenitor’ cell, before becoming a terminally differentiated cell (Fig. 2). Even basic aspects of this model, such as the number of divisions before progression from one compartment to another, are currently experimentally inaccessible and can only be inferred theoretically^38^. In the standard CSCH, only ‘stem cells’ are clonogenic with high probability, but recent modifications have allowed for de-differentiation of TACs into stem cells^22^ and even for terminally differentiated cells (TDs) to ‘move backward’ to become TACs^23^. We propose, instead, that each cell has some mean probability of clonogenicity that is a function of the extent to which a cell is ‘differentiated’, which is, in turn, influenced by microenvironmental variables. We may conceptualize this change in probability of clonogenicity as a cell becomes more ‘differentiated’ as the cell traveling across a non-spatial dimension, which we will call the clonogenic state axis. A cell at the far left of this axis would be clonogenic with high probability (much like a CSC in the canonical model), and a cell at the far right of the axis would be most likely unable to form a colony on its own (a true TD). All other cells would have varying clonogenic potential dependent upon their position on this axis. Accounting for this variance in probability of clonogenicity allows for unification of the original proliferative model and the CSCH and thereby provides an explanation for the dichotomy between sphere-forming ability and true tumorigenicity between ‘marker positive’ cells^39^ and the phenotypic differences observed between spatially separated cells with the same stem marker status^40^.

**Figure 2.**
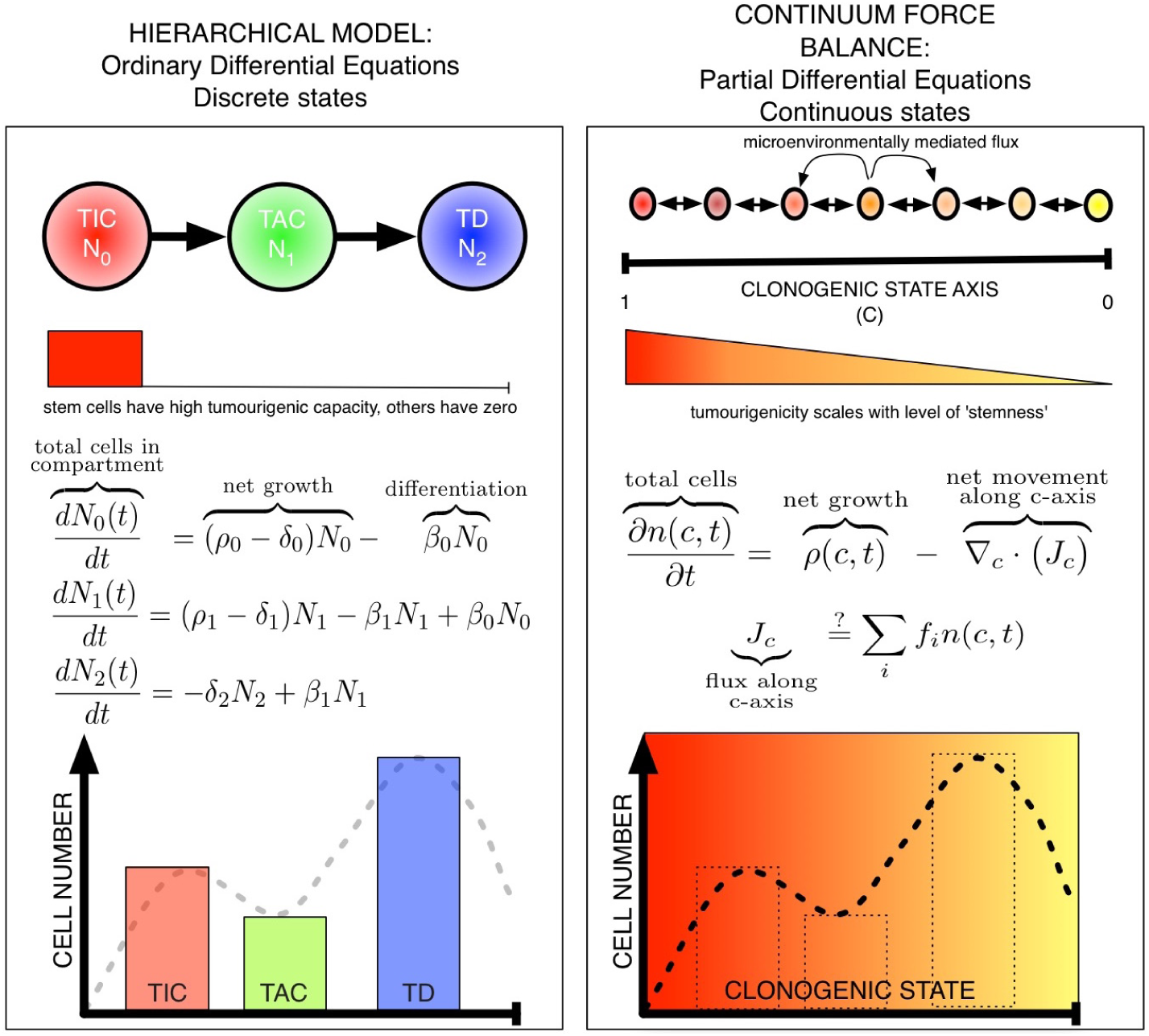
From a discrete, compartment-based description to a continuum. Left: The standard hierarchical stem cell hypothesis, in which each cell type, *i* = (0, 1, 2), self-renews, differentiates and dies at rates *ρ*_*i*_, *β*_*i*_ and *δ*_*i*_ respectively. The given ordinary differential equations thus describe the dynamics of each subpopulation. This rigid, unidirectional model has been extended to various contexts such as dedifferentiation, but the discrete architecture has remained unchanged. Right: The proposed Continuum Force Balance (CFB) model, which allows for a continuum of possible states along the clonogenic state axis (*c*) which could govern a growth rate, *ρ*(*c, t*), incorporates flux of cells along the axis (*J*_*c*_) as driven by microenvironmental ‘forces’ (*f*_*i*_).

**Figure 3.**
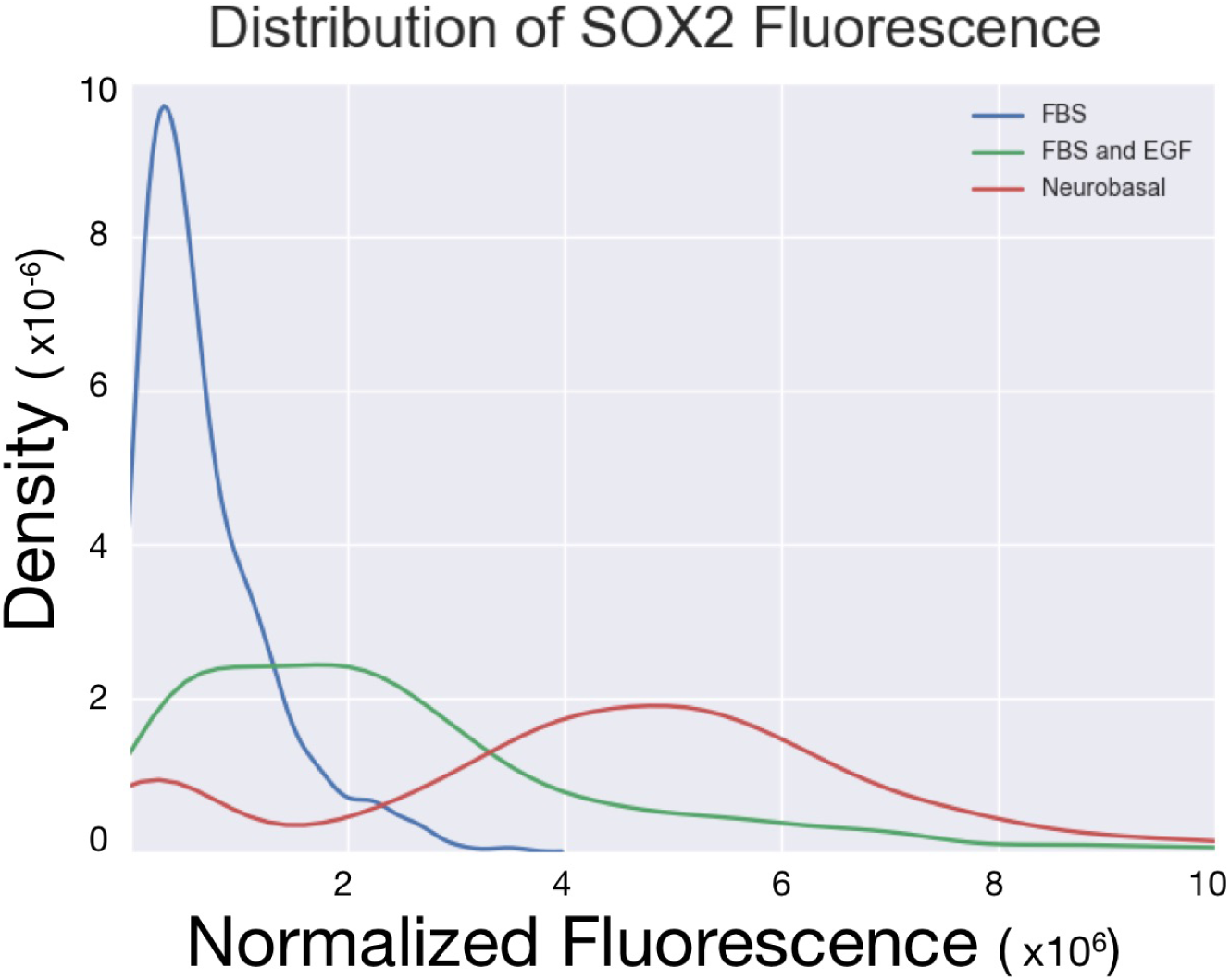
Distribution of SOX2 fluorescence of cell populations grown in differing environments. Analogous to clonogenic state distribution of population of cells, we see a large variation in the population distributions with shifts of the mean of clonogenic potential to the right (corresponding to higher expression of SOX2).

To account for the plasticity that is observed in experiments perturbing the microenvironment^31–33^, we model the microenvironmental variables that affect stemness as ‘forces’ that direct a cell’s movement through the clonogenic state axis. This allows for a single framework in which to view each of these otherwise disparate biological entities, and begs the question as to how these forces are summed in space and time. These forces would, in healthy tissue, be slightly weighted towards the ‘right’ of the axis, such that most cells differentiate as they divide. In a tumor, where there would be a pathologic microenvironment, this balance would be disrupted to the ‘left’ in certain places, acting to skew the cellular population toward classical stemness: an emergent CSC niche.

These different characteristics could account for inter-patient and intra-tumor heterogeneity, and also for the ‘stem cell enrichment’ seen after certain therapies, to include radiation^34^, chemotherapy^10, 35^ and certain microenvironmental factors, such as hypoxia^26^, acidosis^31^, growth factors^36^, and even stromal cell cooperation/cooption^32, 33^ (Table 1). Further, this concept allows for a variable number of differentiation steps and cells of origin^41–43^, dependent on the ‘force balance’ inherent in the environmental context.

**Table 1.** Microenvironmental factors shown to increase stemness in the literature.

| Factor | Tissue type | Source |
| --- | --- | --- |
| Acidosis | Glioblastoma | Hjelmeland et al.<sup>31</sup>, Filatova et al.<sup>44</sup> |
| Hypoxia | Glioblastoma | Conley et al.<sup>26</sup>, Seidel et al.<sup>37</sup>, Soeda et al.<sup>45</sup>, Griguer et al.<sup>46</sup>, Filatova et al.<sup>47</sup>, Kolenda et al.<sup>48</sup> |
| Radiation | Breast and Glioblastoma | Lagadec et al.<sup>25</sup> and Tamura et al.<sup>34</sup> |
| Chemotherapy | Glioblastoma and Liver | Chen et al.<sup>10</sup> and Hu et al.<sup>35</sup> |
| EGF | Brain | Doetsch et al.<sup>36</sup> |
| HGF and Wnt | Colon | Vermeulen et al.<sup>32</sup> |
| IL-6 and CXCL-7 | Breast | Liu et al.<sup>33</sup> |

### Quantifying the effect of microenvironmental variables: a multidisciplinary task

The evolving CSCH has been driven by the growing body of literature suggesting that microenvironmental signals can affect stemness (Table 1). We seek to coalesce these signals into a single ‘force’ term that will enable dynamic modeling of a spatially heterogeneous hierarchically organized tumor with a continuum approximation. To do this in a way that is meaningful however, will require adoption of this concept by experimental as well as theoretical scientists. Experiments which currently show the qualitative effect of microenvironmental perturbations on ‘stemness’ must be done quantitatively, and a standard measure of this ‘force’ will have to be ascribed. This measure of force must be descriptive enough to identify the change in the distribution of a population of tumor cells from its initial state to a later state along the clonogenic state axis, and the time over which the change occurred.

To coalesce the individual perturbations into a single measure, quantitative experiments focussing on dose-response relationships using different ‘stem’ markers must be undertaken and scaling laws defined. Once this is accomplished, experiments focussing on synergy between different factors can begin, and as each of the listed microenvironmental perturbations identified so far is measured in different ways, the need for conversion factors arises. While understanding all possible interactions would require many different combinations of the factors, even a minimal set of baseline quantitative experiments would shed light on what is now only a qualitative understanding of the underlying biology. Once these baseline measurements have been accomplished, the proposed model, along with continuum models of the microenvironment (e.g. Anderson et al.^49^), could be used as a predictive tool to understand the temporal evolution of a clonogenic distribution and how it relates to factors such as invasion, heterogeneity and treatment resistance.

To highlight the manner in which perturbations affect the distributions of clonogenic cells, experiments involving CD133 positive cells from a patient-derived xenograft cell line were assayed for SOX2 expression, as described in the Methods section. SOX2 is a transcription factor involved in the maintenance of the pluripotency of embryonic stem cells, and in the experiments we present, serves as a marker of clonogenicity, with higher expression per cell corresponding to a ‘left’ shift on the clonogenic state axis. Prior work has shown that silencing of SOX2 results in a significant decrease in clonogenicity, underscoring its role as a critical measure of clonogenicity in a cell^50^.

In the experiments performed, we consider three different conditions of cell treatment, each of which shifts the distribution of clonogenicity in unique ways. Cells are treated with either fetal bovine serum (FBS), a known differentiating agent for cancer stem cells, epidermal growth factor (EGF) and FBS, or neurobasal media, with EGF and fibroblast growth factor (FGF). EGF, as reported in Table 1 has been shown to induce stemness in neural precursors, and it is also thought that neurobasal media (the combination of FGF and EGF) should increase stemness as well, though we show that, in fact, these shift the distribution of clonogenicity in distinct ways.

This dataset demonstrates how each force perturbs the distribution of cells over the clonogenic state axis in unique ways, and how the forces are unique, in that they cannot be summed independently to obtain a combination distribution. Further, we note that *all* cells at the beginning of the experiment were CD133 biomarker positive, suggesting that the gradient in clonogenicity as elucidated by SOX2 expression varies significantly among this population. Also, this suggests that binary classification by CD133 expression alone does not provide a significant degree of resolution of the clonal capabilities of the cell population. In this vein, it is critical to note that the distribution of clonogenicity acts as a hidden parameter in experimental data. That is, it is not directly observable with particular cellular surface markers used in a binary manner, and in this way, may explain inconsistencies between populations of stem cells behaving vastly differently, despite being similar in terms of biomarker positive proportion. In this sense, it may be that a biomarker selects only for some subset of the distribution of clonogenic cells, but ignoring the intermediate clonogenicity of the cells not selected by the biomarker may give rise to the discrepancies between the expected dynamics and the experimental observations.

### Ramifications for carcinogenesis and progression: from a static to dynamic understanding

As we continue to understand and classify the factors that exert these putative ‘forces’ which change the distribution of cells along the clonogenic state axis, we will be better able to understand how the dysregulation of the force balance affects not only tumor progression, but also carcinogenesis. For instance, it may be that pre-cancerous lesions are more ‘left-skewed’ than their healthy counterparts, and that cancer-associated stroma is reacting in a physiologic manner to pathologic signals (‘left forces’). Further, a loss of structure of distribution along the clonogenic axis, secondary to a perturbation or imbalance of the cellular ‘forces’, may define a cancerous tissue, or one which is changing from pre-malignant to malignant. Such a situation could be reached by multiple combinations of genetic mutations or environmental perturbations in both the pre-malignant and healthy associated tissue nearby. This provides a unification of the context-specific hypotheses of cancer that have plagued the CSCH for so long, in which clonogenic cancer cells mixed with healthy tissue, may be affected by a physiologic ‘right force’, and revert to a healthy state^51^.

By reformulating the CSCH with our continuum force balance (CFB) model, expressed mathematically as the partial differential equation (PDE) in Fig. 2, we are also presented with novel opportunities to rethink the utility of cell surface markers previously attributed to ‘stemness’. We posit that the degree of positivity of a particular marker (applied in a binary fashion) gives a measure of the proportion of the tumor population that exists to the left of a threshold on the clonogenic state axis. In this sense, one may hypothesize that the use of thresholds from several markers in combination may give a strong sense of the underlying distribution of the tumor cells along this axis. We argue that this distribution of clonogenicity is a more robust measure of a tumor’s state, unmasking underlying complexity, than simply a proportion of CSCs with a single binary marker. Importantly, this may provide the necessary explanation for the differences observed when reconstituting tumor populations from ‘purified’ populations of biomarker-positive cell populations.

### Spatial heterogeneity allows for the stem cell niche as an emergent phenomenon

To this point, we have offered a mathematical construct which describes the distribution of cancer cells along a continuous ‘clonogenic probability’ axis by means of a ‘clonogenic force’ based model, which we submit should replace the hierarchical CSCH. We have described ways in which this new construct can better explain the existing biological observations and also ways that it can open the field to new lines of questioning. Specifically, we have suggested novel methods to 1) quantitatively define the effects of microenvironmental perturbations, 2) characterize the makeup of a tumor by utilizing suites of cell surface markers, and 3) represent the effect of extrinsic microenvironmental pressures or genetic alterations as ‘forces’ along a non-spatial continuum axis.

We emphasize that the equation described in Figure 2 (right) provides a concrete illustration of a more general framework within which a continuum of clonogenicities may be incorporated. To better model the reality of spatial heterogeneity observed in solid tumors, it is straight forward to extend this formulation to include physical space (**x**), thus,

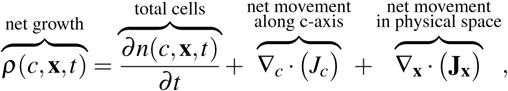

where *J*_*c*_ could be equal to *n*(*c,* **x***, t*) *f* (*c,* **x**) or another functional form, and **J_x_** can be represented by any appropriate function modeling cell motility^52^. This addition allows for varying functions for birth, death and ‘forces’, both as the cell differentiates (as a function of *c*), as a function of microenvironmental factors which would vary by physical location, **x**. The possibilities for different functional forms for these fluxes (*J*_*c*_ and *J***_x_**) represents a rich field for both theoretical and experimental work. As written, this formulation can now represent the stem cell niche as an emergent phenomenon secondary to microenvironmental conditions and cellular characteristics; not unlike the current understanding of the hematopoietic stem cell niche^53^. More importantly, we now offer a single mechanism to explain how different types of cancer stem cell niches could be created and maintained (e.g. the perivascular^54^, invasive^55^, and hypoxic^37^ niches in glioblastoma). Further, describing the tumor as a distribution of cells with varying ‘clonogenicity’ and state-specific replication and differentation rates, rather than arbitrarily discretised compartments, we are able to model the changing nature of a tumor over time and space in a new, quantitative, way.

## Conclusion

The current state of modeling the CSCH has reached a point where there are a number of biological observations that challenge assumptions in previous models, such as a continuity of stemness, plasticity of the stem phenotype and significant effects of the physical microenvironment on stemness. To remedy this, we suggest revising this restrictive structure of the standard hierarchy and offer instead a continuum force balance model, which allows for new quantitative observations to be interpreted in terms of the clonogenic state force within a consistent mechanistic framework. This novel formulation serves to settle the dissonance between the proliferative hypothesis and the CSCH and provides a single, integrated framework by which to capture several puzzling phenomena in cancer biology. This description provides new opportunities for both theoretical and experimental insights within the field of CSC research, which has presented many results but, to date, frustratingly few translatable insights.

## Supplementary Information

### Experimental Methods

#### Glioblastoma stem cell preparation and maintenance in culture

Human glioblastoma stem cells were enriched by CD133 Macs Beads (Miltenyi Biotech, San Diego, CA) following patient-derived xenograft tumor dissociation using the Papain Dissociation System (Worthington Biocheical Co., Lakewood, NJ) as described previously^56^. They were maintained as sphere cultures in Neurobasal medium supplemented with B27, L-glutamine, sodium pyruvate, penicillin, streptomycin (Thermofisher, Waltham, MA), EGF and basic FGF (each at 20 ng/ml, R&D Systems, Minneapolis, MN).

#### Quantification of Sox2 expression

To quantify Sox2 nuclear expression levels in individual cells, we used a quantitative immunofluorescence approach as described previously^57^. To establish cancer stem cell monolayer cultures for immunostaining, coverslip (22 × 22 mm, Thickness 1.5, Corning, NY) were coated with Geltrex (Thermofisher, Waltham, MA) as described previously^58^. Geltrex, which is rich in laminin, was used as a cell attachment substrate to provide a stem-cell-supporting environment^59^. Glioblastoma CSC spheres were dispersed to single cells with Accutase (Bioledgend, San Diego, CA) and plated on to Geltrex coated coverslips in 6 well plates with CSC maintenance medium (200,000 cells per well). After 3 days of culturing since changing the medium to one that contained either fetal bovine serum (FBS, 10%; Sigma-Aldrich, St. Louis, MO), FBS & EGF (20 ng/ml), or EGF & FGF (20 ng/ml for each), the cells were fixed with 4% paraformaldehyde. Following permeabilization and blocking with 0.1% (w/v) Triton X-100, 2% (v/v) normal donkey serum containing PBS, monolayers were incubated with a specific antibody against Sox2 (1:500 dilution, mouse monoclonal (clone #245610) anti Sox2 antibody, R&D Systems, Minneapolis, MN). Donkey anti mouse IgG conjugated with Cy3 (Jackson ImmunoResearch, West Grove, PA) was used to detect bound anti Sox2 antibody, and DNA was stained with Hoechst 33342 (Polysciences Inc., Warrington, PA). Digital images of Sox2 and DNA stainings of the same fields were taken using a fluorescent microscope (Leica DM5000B) equipped with a digital camera (Leica DFC310FX).

Quantitative image analysis was performed using ImageJ^57^. Sox2 expression levels in individual nuclei were quantified by integrating the pixel intensity values of Sox2 staining in the nuclear regions, which were defined by Hoechst staining.

